## Supplementary Table 1 for "Rtf1-dependent transcriptional pausing regulates cardiogenesis"

**Supplementary Table 1.** Sequences of primers used for quantitative real-time PCR analysis of gene expression in mouse embryonic stem cells.

| Gene | Forward Primer Sequence (5' to 3') | Reverse Primer Sequence (5' to 3') |
|---|---|---|
| ***Ppia* (control)** | GTCTCCTTCGAGCTGTTTGC | GATGCCAGGACCTGTATGCT |
| ***Brachyury*** | TGTGAGAGGTACCCAGCTCTAAGGAA | TATCATGGGACTGCAGCATGGACA |
| ***Ncam1*** | CATCATCTGGAAACACAAAGGCCGAG | GATCTGCAGGTAGTTGTTGGACAGGA |
| ***Afp*** | TGTCTGCAGGATGGGGAAAAAG | GCCATTCTCTGCGTGAATTATG |
| ***Nkx2-5*** | TGCGCTCACACCCACGCCTTTCTCA | CCAGACGCCAGGCTACGCTGCT |
| ***Myh6*** | CACTCAATGAGACGGTGGTG | GTGGGTGGTCTTCAGGTTTG |
| ***Nppa*** | TACAGTGCGGTGTCCAACACAGAT | TGACCTCATCTTCTACCGGCATCTTC |
