## Supplementary Table 2 for "Rtf1-dependent transcriptional pausing regulates cardiogenesis"

**Supplementary Table 2.** Predicted identities of cell types belonging to each Seurat cluster identified in analysis of 11-12 somite stage control and *rtf1* morphant embryo integrated multimodal (ATAC + Gene Expression) single cell sequencing datasets. Predicted identities were based on manual inspection of marker gene expression.

| Seurat Cluster | Predicted Cell Type |
|---|---|
| **0** | Brain/CNS |
| **1** | Posterior Segmental Mesoderm |
| **2** | Hatching Gland cluster 1 |
| **3** | Periderm cluster 1 |
| **4** | Basal Cells cluster 1 |
| **5** | Notochord cluster 1 |
| **6** | Pre-somitic Paraxial Mesoderm |
| **7** | Periderm cluster 2 |
| **8** | Basal Cells cluster 2 |
| **9** | Lateral Plate Mesoderm |
| **10** | Forebrain |
| **11** | Neural tube/Neurons |
| **12** | Adaxial cells |
| **13** | Caudal Blood Island Precursors |
| **14** | Pharyngeal Mesenchyme |
| **15** | Optic Vesicle |
| **16** | Midbrain-hindbrain Boundary |
| **17** | Fast Muscle Precursors |
| **18** | Floorplate |
| **19** | Endothelial (high *flt4*) |
| **20** | Otic Placode |
| **21** | Pronephric Mesoderm |
| **22** | Pre-cardiac Mesoderm |
| **23** | Neural Crest |
| **24** | Somite cluster 1 |
| **25** | Endoderm |
| **26** | Rostral Blood Island Precursors |
| **27** | Ionocyte / Mucus Cell Precursors |
| **28** | Epidermis |
| **29** | Neurons cluster 1 |
| **30** | Endothelial (low *flt4*) |
| **31** | Notochord cluster 2 |
| **32** | Neurons cluster 2 |
| **33** | Notochord cluster 3 |
| **34** | Yolk Syncytial Layer |
| **35** | Notochord cluster 4 |
| **36** | Hatching Gland cluster 2 |
| **37** | Notochord cluster 5 |
| **38** | Somite cluster 2 |
